# Differential Principal Components Reveal Patterns of Differentiation in Case/Control Studies

**DOI:** 10.1101/545798

**Authors:** Benjamin J. Lengerich, Eric P. Xing

**Affiliations:** Computer Science Department, Carnegie Mellon University, Pittsburgh, PA 15213; Machine Learning Department, Carnegie Mellon University, Pittsburgh, PA 15213; Petuum Inc., Pittsburgh, PA 15222

## Abstract

Dimensionality reduction is an important task in bioinformatics studies. Common unsupervised methods like principal components analysis (PCA) extract axes of variation that are high-variance but do not necessarily differentiate experimental conditions. Methods of supervised discriminant analysis such as partial least squares (PLS-DA) effectively separate conditions, but are hamstrung by inflexibility and overfit to sample labels. We would like a simple method which repurposes the rich literature of component estimation for supervised dimensionality reduction.

We propose to address this problem by estimating principal components from a set of difference vectors rather than from the samples. Our method directly utilizes the PCA algorithm as a module, so we can incorporate any PCA variant for improved components estimation. Specifically, Robust PCA, which ameliorates the deleterious effects of noisy samples, improves recovery of components in this framework. We name the resulting method *Differential Robust PCA* (drPCA). We apply drPCA to several cancer gene expression datasets and find that it more accurately summarizes oncogenic processes than do standard methods such as PCA and PLS-DA. A Python implementation of drPCA and Jupyter notebooks to reproduce experimental results are available at www.github.com/blengerich/drPCA.

## Introduction

Many bioinformatics datasets contain more features than samples, often because -omic assays profile many biomarkers but collecting data from a large number of individuals is costly. This reduces the statistical power of machine learning algorithms to distinguish signal from noise, a problem known as the curse of dimensionality (1). One way to alleviate this problem is to reduce the number of features in the dataset.

Principal components analysis (PCA) is the most popular way to summarize high-dimensional datasets. PCA projects datapoints onto the axes of major variation (2). While PCA minimizes reconstruction error of the training data as measured in Euclidean distance, the selected axes (and the resulting data representations) are not guaranteed to be biologically meaningful. For example, in gene expression studies, the axes of major variation often correspond to technical artifacts or biological processes which are not tightly regulated (3). These high-variance process are selected by PCA in order to reduce recovery error but they may not efficiently characterize the phenomenon of interest. Projecting data onto the top principal components can thus discard valuable information about tightly-regulated biological processes.

We propose to learn the principal components which summarize the *differences* between groups rather than optimize reconstruction error. If the low-dimensional representations succinctly capture the variation between groups, they may be more useful for understanding the processes of differentiation.

To estimate components of differentiation, we apply PCA-based methods to a set of vectors that define the difference between case and control groups. We call this framework *Differential PCA* (dPCA) and find that *Differential Robust PCA* (drPCA) compares favorably to supervised dimensionality reduction techniques while maintaining simplicity, modularity, and extensibility. Beyond the improved performance on the datasets presented in this article, we are excited about the possibility of the framework to be expanded by incorporating other techniques of dimensionality reduction techniques that have been developed for biological datsets.

A Python implementation of drPCA, as well as Jupyter notebooks to reproduce experimental results, can be downloaded from ^1^.

## Motivating Example

Shown in Figure 1 is an illustration of various dimensionality reduction methods on a toy dataset. Fig. 1a depicts the twodimensional datapoints we are interested in compressing. This dataset has two different clu ers: a backgrou dataset (orange) generated by 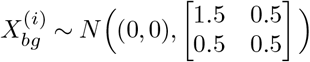, and a foreground dataset orange) generated by th ample-specific process 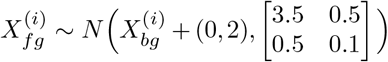. Each foreground datapoint is thus vertically offset from a background setting and perturbed by Gaussian noise.

When analyzing this dataset by components analysis, there are many experimental questions that may be asked. To understand the differences between foreground and background data, we may want to identify the vertical axis as the component of differentiation.

**Fig. 1.**
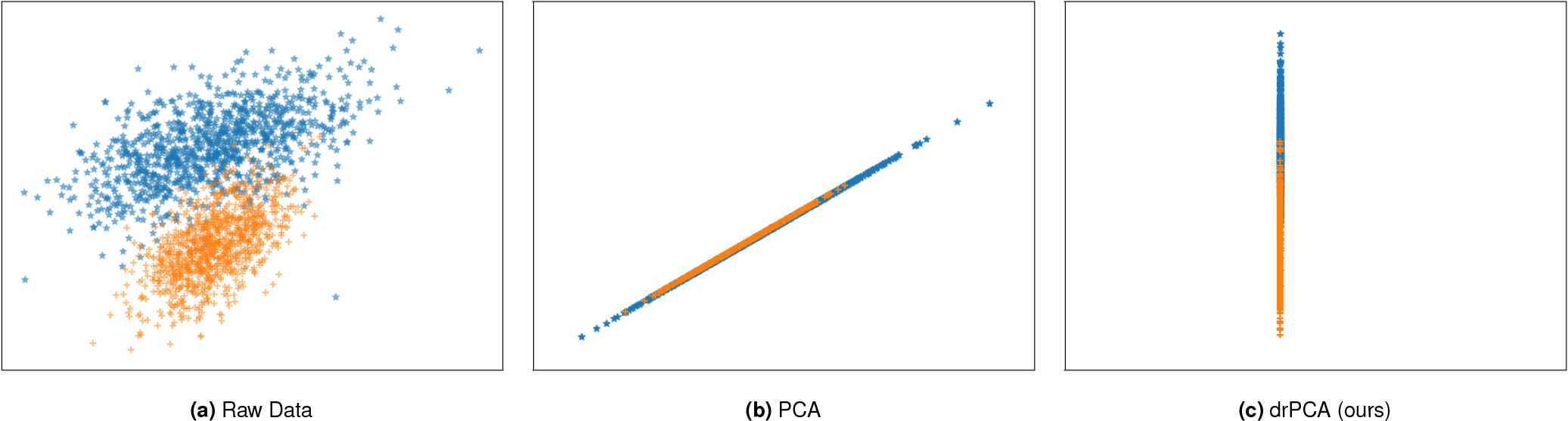
A toy example of dimensionality reduction. (a) Foreground datapoints (blue) are vertically offset from the background datapoints (orange) and perturbed by zero-mean noise. (b) Projection of datasets onto the first component recovered by PCA. (c) Projection of datasets onto the first component recovered by drPCA. In this setting, drPCA successfully recovers the axis which differentiates the foreground and background data, while PCA projects both groups onto an axis which does not distinguish foreground samples from background samples.

PCA projects the datapoints onto the axis of major variance (Fig. 1b), but this axis does not distinguish foreground and background samples. As a result, the clusters are completely overlapping. In contrast, drPCA identifies the axis which differentiates foregrond and background samples (Fig. 1c).

## Differential PCA

A differential PCA method (dPCA) was first propsoed by (4) as a targeted solution for ChIP-Seq data in which the goal is to identify protein binding capacities which differentiate experimental settings. Because Ji *et al.* specifically formulated dPCA for ChIP-Seq experiments, they make the natural assumption that there are many different experimental settings and many replicates of each setting. As a result, their proposed method first computes the mean of the data samples in each group, and then performs PCA on the dataset of differences between means. This process leads to selection of axes which characterize the protein-binding process studied in CHiP-Seq data, and has been successfully applied in several studies (5, 6).

When applying this idea to settings beyond ChIP-seq data, we encounter a major problem: the number of principal components is strictly less than the number of datapoints. If each datapoint is defined as the mean of the replicates for an experimental setting, then we must always have more experimental settings than desired components. For the common task of case/control data, no principal components are defined, and only a single prinicpal component can be extracted if the dataset is “grounded” by adding a zero vector.

To alleviate this problem, we do not take the means of each cluster. Instead, our dataset of difference vectors is calculated directly from pairs of samples. Given a list of matched samples, we calculate the high-dimensional difference vectors between the samples in the foreground set and the samples in the background set. To make this simple methodological change explicit, we will refer to the method of Ji *et al.* as dPCA-Mean, and the non-averaged method as dPCA.

Running components analysis directly on this differential dataset would identify axes of variance in the differences, but we are seeking to summarize the differential vectors. To analyze the differences as vectors, we “ground” the differential dataset by adding pseudo-samples of zero. With the number of zeros equivalent to the number of difference vectors, the principal components of the difference dataset summarize axes of differentiation between the sample clusters. This procedure is summarized in Alg. 1.

This construction allows us to utilize the well-studied suite of denoising and structured variants of PCA by replacing the call to *PCA* in line 6 of Alg. 1 with a call to a variant of PCA.

### Algorithm 1 Differential PCA

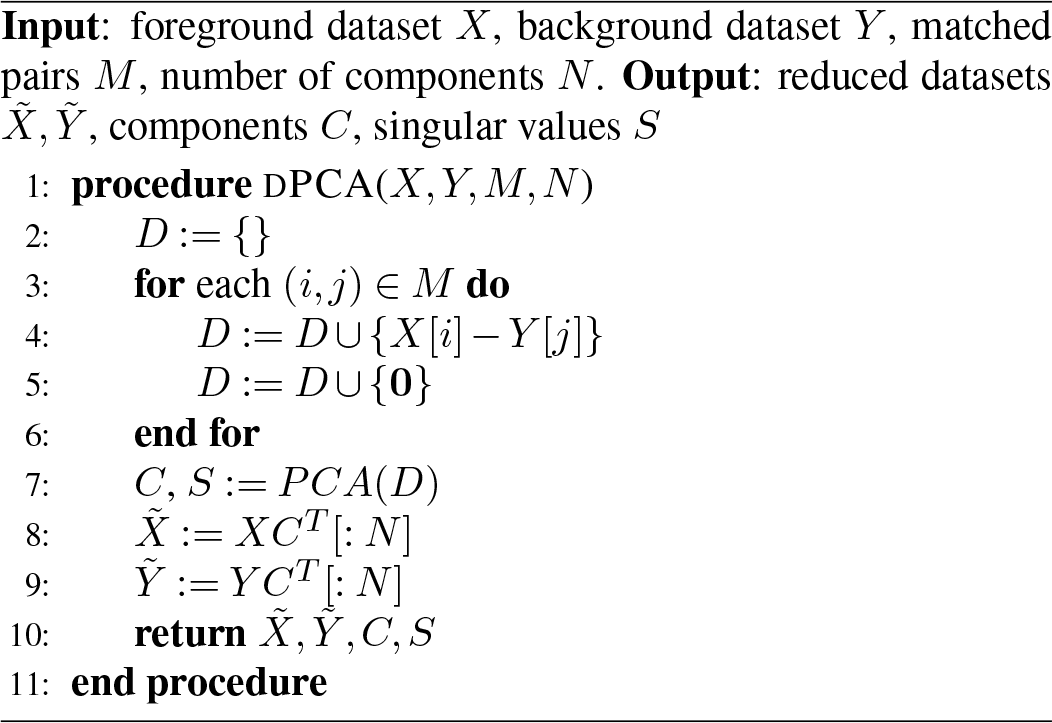

### Differential Robust PCA

As described below, Robust PCA (rPCA) decomposes the data into the sum of a low-rank component and a sparse component to increase stability in the presence of noise (7). We can incorporate rPCA in the dPCA framework by replacing the function call to *PCA* in line 6 of Alg. 1 with a call to *rPCA*. We call this method Differential Robust PCA (drPCA). In our experiments, we see that drPCA performs extremely well in differential datasets, even when case and control samples are not from matched sources. This highlights the benefit of the dPCA framework to reuse the rich literature of methods for components analysis.

### Unmatched Samples

If samples are not taken from matched individuals, it becomes necessary to construct a new dataset of matched pairs. In this case, we can generate a matched dataset by uniformly selecting datapoints from each condition to be matched. This process produces a noisy differential dataset which is handled by the denoising aspects of drPCA.

## Related Work

The most popular methods for dimensionality reduction are unsupervised linear methods, which find a linear transformation that projects the high-dimensional data points onto a nearby low-dimensional subspace. Of these, the most widely-known method is principal component analysis (PCA), which learns a set of linearly orthogonal features that represent the directions of maximal variance in the original data (8). Since PCA was first introduced, many variants have been developed. For example, Sparse PCA (9) uses an elastic net penalty to encourage element-wise sparsity in the projection matrix, while Independent Component Analysis (10) (ICA) recovers statistically independent components to separate signal sources. Robust PCA (rPCA) learns to decompose the data into the sum of a low-rank component and a sparse component, leading to increased stability in the presence of noise (7, 11).

There are also approaches that use richer models for the underlying latent representation of the data. These include methods that perform simultaneous dimensionality reduction and feature selection (12) or non-linear dimensionality reduction methods. Among these deep models, unsupervised methods such as variational (13) and denoising (14) autoencoders seek to learn latent features by optimizing data reconstruction in a bottlenecked architecture. These methods may also be extended to the supervised case (15); however, the deep architecture often requires more samples than are available for high-dimensional genomic assays. Additionally, it can be difficult to analyze the non-linear compression functions in a sample-agnostic way. For these reasons, we consider only linear dimensionality reduction techniques in the remainder of this paper.

### Supervised Dimensionality Reduction

Supervised methods use sample labels in order to produce more meaningful data representations. Here, we describe several supervised methods frequently used in bioinformatics analyses.

#### Linear Discriminant Analysis

Linear Discriminant Analysis (LDA) (16) seeks to separate datapoints according to sample labels. To do so, LDA analytically maximizes cluster separation under a linear model. As a result, LDA produces clusters which are extremely well-separated. However, because LDA uses a closed-form solution, it can be difficult to extend the framework to recover desired structure in the components and can make it challenging to recover biologically interpretable components.

#### Supervised PCA

Supervised PCA (17) modifies traditional PCA by considering only the subset of explanatory variables which have sufficient correlation with the sample labels. In this way, Supervised PCA estimates sparse components which contain only features that may be predictive of the phenomenon of interest. However, as combinations of these features, the components may not describe the differences between sample groups.

#### Partial Least Squares and Canonical Correlation Analysis

Partial least squares (PLS) (18) and Canonical Correlation Analysis (CCA) (19) are bilinear factor models which fit linear projections for both outcomes and regressors. If categorical outcomes are used, as in sample labels, PLS is called PLS-Discriminant Analysis (PLS-DA) (20). While PLS-DA has many pleasing qualities, including handling high dimensions and multicolinearity well (for which CCA struggles), the method can be difficult to extend. Specifically, we would like to incorporate biological knowledge into our component estimation procedure, but it is not immediately clear how to modify PLS-DA to achieve this without implementing a new optimization procedure. This motivates us to consider the simple framework of differential PCA, in which well-tuned PCA variants may be used interchangably.

#### Contrastive PCA

A recently developed method for case/control data is Contrastive PCA (cPCA) (21). cPCA identifies axes that have large variance in the foreground samples but small variance in the background samples. In this way, cPCA can identify patterns of differentiation in the diseased state, potentially implicating dysregulated pathways. As shown in Section, cPCA and drPCA tend to select different components from cancer gene expression data; this suggests that the patterns which most differentiate cancers from one another are not the oncogenic processes which caused the cancers.

#### Differential Expression

For gene expression data, a closely related framework is differential expression (DE). In DE analyses, statistical tests are used to assess the probability that the means of the expression amounts for each gene are the same in both experimental settings. While both DE and dPCA seek to analyze the differences between two experimental conditions, there are stark differences. Firstly, dPCA performs subspace mapping rather than feature selection. In addition, we can induce structure in the recovered components via Bayesian methods, but this would be difficult in DE studies due to the univariate testing nature.

#### Discriminant Analysis of Principal Components

For SNP data, Discriminant Analysis of Principal Components (DAPC) is popular for identifying and describing clusters of genetically related individuals (22). However, DAPC uses population structure data that is specific to SNP assays. In contrast, our proposed method is relevant to any case/control study.

## Experiments

To understand the behavior of these dimensionality reduction methods, we perform several experiments. First, we use simulated data to quantify how well drPCA recovers axes of differentiation from data. Next, we turn to cancer gene expression analysis to inspect the biological processes which differentiate tumor samples from healthy controls.

### Simulated Data

We simulate data according to a mixture of two clusters. The background dataset is generated by 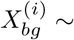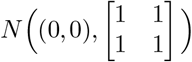, and a foreground dataset is generated by the sample-specific process 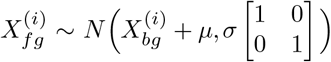, creating an offset of *μ* between the foreground and background clusters. In this experiment, each datapoint consists of 1000 dimensions. To simulate biological data in which many features are unrelated to the process under study, we make the offset *μ* sparse, with only 5 non-zero entries. We generate *n* foreground and background datapoints according to this structure, and measure the cosine similarity between the estimated axes of differentiation and *μ*. Results for various *n* are shown in Figure 2.

**Fig. 2.**
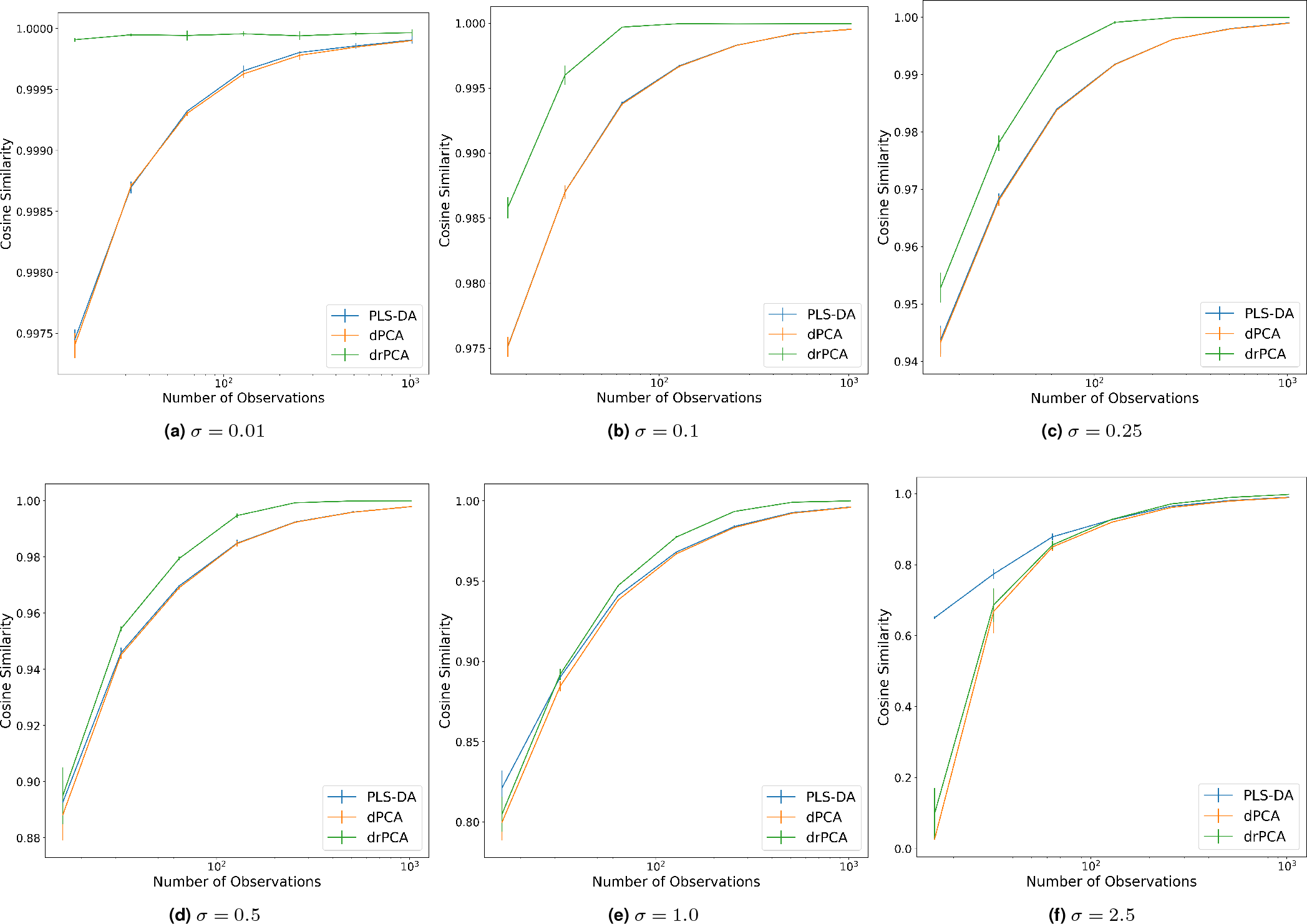
Recovery of axes of differentiation from simulated data for varying levels of noise governed by *σ*. On the y-axis, we show the cosine similarity between the true axis of differentiation and each method’s first component. Results are averaged over 5 experimental settings, with standard deviations depicted by error bars. Results from other baseline methods are omitted because they all have cosine similarity below 0.8. In settings with low noise and moderate sample sizes, drPCA outperforms PLS-DA. In settings with large noise or very few samples, PLS-DA outperforms drPCA.

As shown in Figure 2, drPCA outperforms the baselines at recovering *μ*, the axis of differentiation, in settings with low noise and moderate sample sizes. In settings with large noise or very few samples, PLS-DA outperforms drPCA. Other baseline methods have extremely poor performance on this task.

### Cancer Gene Expression Studies

We investigate several RNA-seq gene expression datasets from The Cancer Genome Atlas^2^. These datasets profile cancer patients and contain tumor samples with some matched healthy tissues. We inspect three different cancer types: Breast Invasive Carcinoma (BRCA), Lung Adenocarcinoma (LUAD), and Glioblastoma Multiforme (GBM). These datasets contain 1102, 533, and 312 samples from cancer tissues, respectively. In addition, they contain 113, 59, and 10 samples from control tissues, respectively. The datasets are extremely high-dimensional; the datasets contain 15584, 14533, and 30584 distinct transcripts after thresholding features for a minimum standard deviation. In addition to these disease-specific datasets, we also evaluate dimensionality reduction on the combination of the three datasets (Combined). For the GBM dataset, we supplement the matched differences with 750 unmatched differences produced by randomly matching case and control samples. To evaluate the performance of the estimated components, we hold out 40% of the patients from each dataset for downstream tasks.

### Differential Components Separate Case and Control Samples

We first visually inspect the clusters induced in the top two components of each method. The projected data for the Combined dataset are shown in Fig. 3; similar results for the BRCA, LUAD, and GBM datasets are available at github.com/blengerich/drpca.

Standard PCA (Fig. 3a) and unsupervised variants (Fig. 3b, Fig. 3c) project the data onto axes which do not differentiate case and control samples. While these axes may be useful for characterization of the samples, they are unlikely to correspond to processes which are causal for the tumors. In addition, we see that Contrastive PCA (Fig. 3d) effectively identifies processes which have high variance in the cancer samples but low variance in the control samples. These components may correspond to processes which are dysregulated in tumors but not oncogenic, a hypothesis supported by inspection of the component loadings (Tab. 1).

**Fig. 3.**
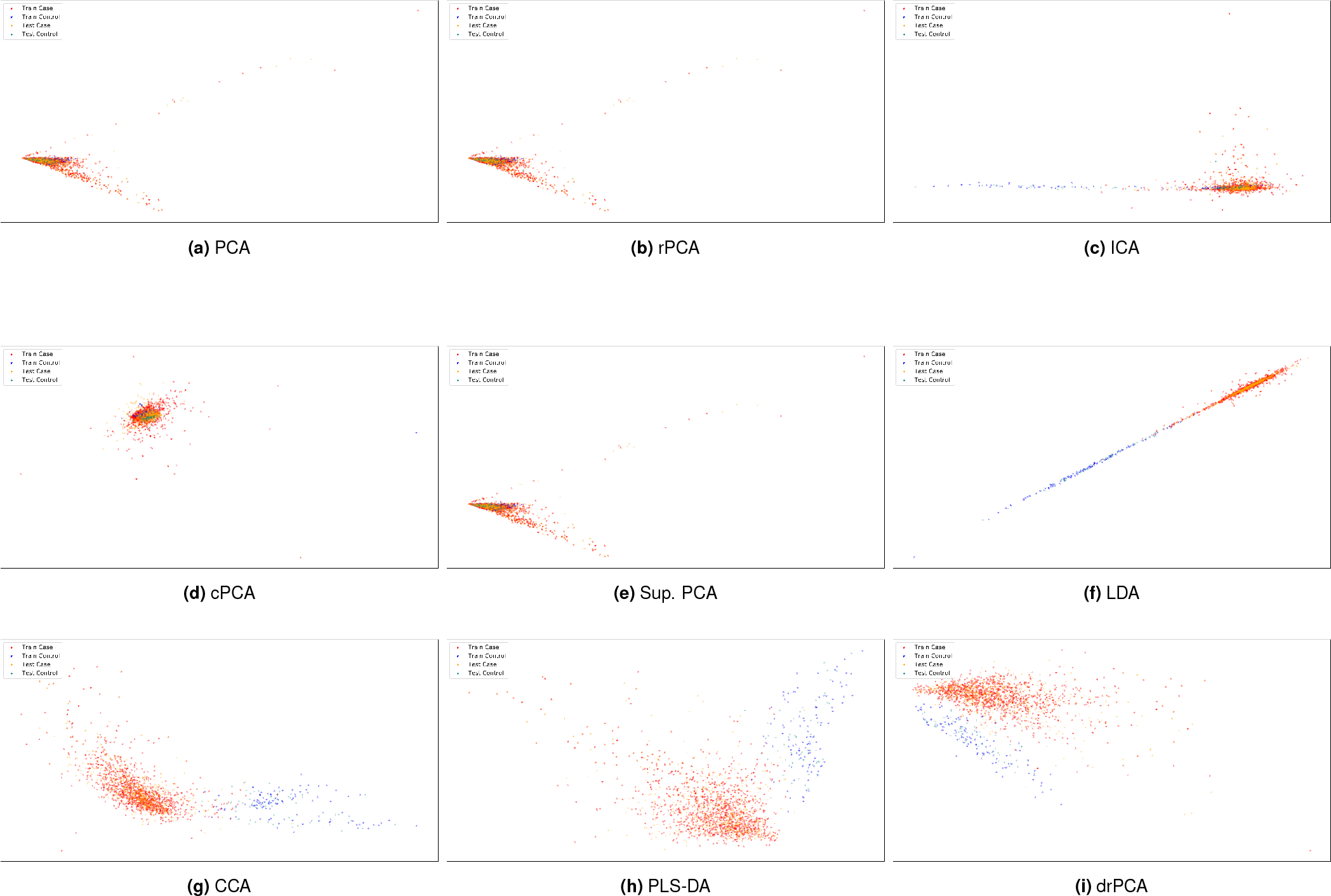
Low-dimensional representations of samples from the combined cancer dataset. The supervised methods LDA (f), CCA (g), PLS-DA (h), and drPCA (i) all separate case and control samples.

In contrast to the unsupervised dimensionality reduction methods, the supervised methods (Figs. 3e,3f,3g,3h,3i) all effectively separate case and control samples, with separability transferring betwen the training and test sets. Of these, drPCA and PLS-DA produce the most visually distinctive clusters, with mean silhouette scores greater than 0.6.

### Differential Representations are Useful for Predictive Tasks

To test the presence of biological signal in these low-dimensional representations, we measure the performance of random forest (RF) classifiers for two tasks. First, we train the RFs to label case and control samples. After learning components from the training set, we optimize a RF classifier to predict case/control labels by cross-validation on the same training set. Plotted in Fig. 4 are the performances of the classifiers on the held-out test set projected into a given number of components. As expected, the supervised methods CCA, PLS-DA, LDA, dPCA, and drPCA all significantly outperform the unsupervised methods because they use the sample labels. However, the scientific utility of the representations produced by PLS-DA, LDA, and CCA are questionable; changing the task severely degrades predictive performance.

After reducing dimensionality based on case/control labels, we train another RF to predict the tissue of origin for each sample (recall that the combined dataset is composed of samples from three different tissue types). As shown in Fig. 5, the LDA representations contain very little information that is predictive of this task. As a result, the AUROCs using this method does not surpass 0.6. The representations from CCA and PLS-DA also struggle on this simple task. In contrast, the representations from dPCA and drPCA are among the best-performing representations for this task. This suggests that the differential components are biologically meaningful while the baseline supervised methods overfit to the labels.

### Differential Components Summarize Oncogenic Patterns

Do the components which separate tumor samples from control samples give high weight to oncogenic processes? To answer this question, we sort the variables according to the magnitude of the weight in the first component of each method.

**Fig. 4.**
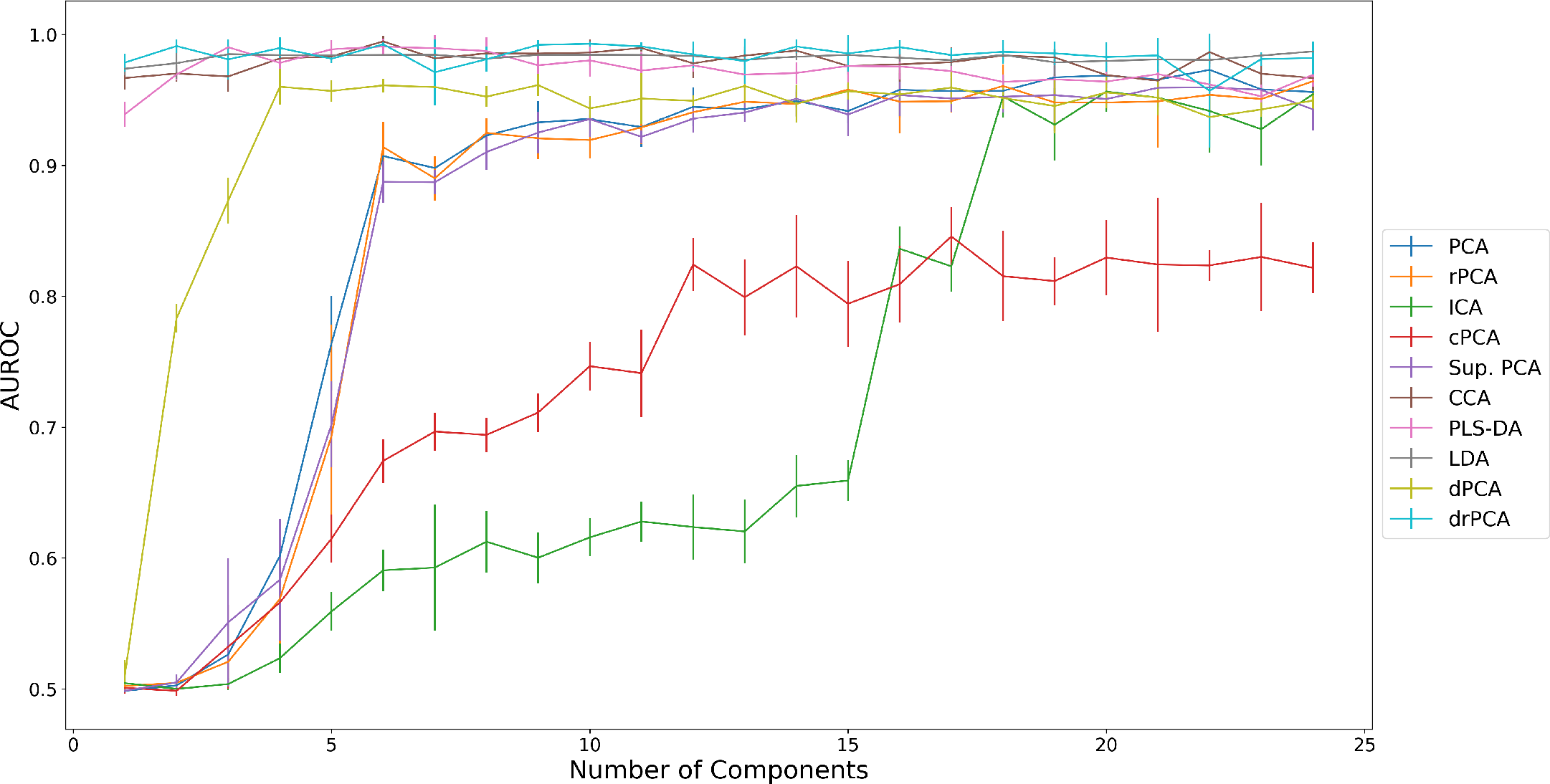
Prediction of case/control status from the Combined dataset. The y-axis is the areas under the Receiver Operating Characteristic curves (AUROCs) of the prediction, with x-axis indicating the number of components used in the representations. Errorbars indicate the standard deviation over 3 train/test splits.

**Fig. 5.**
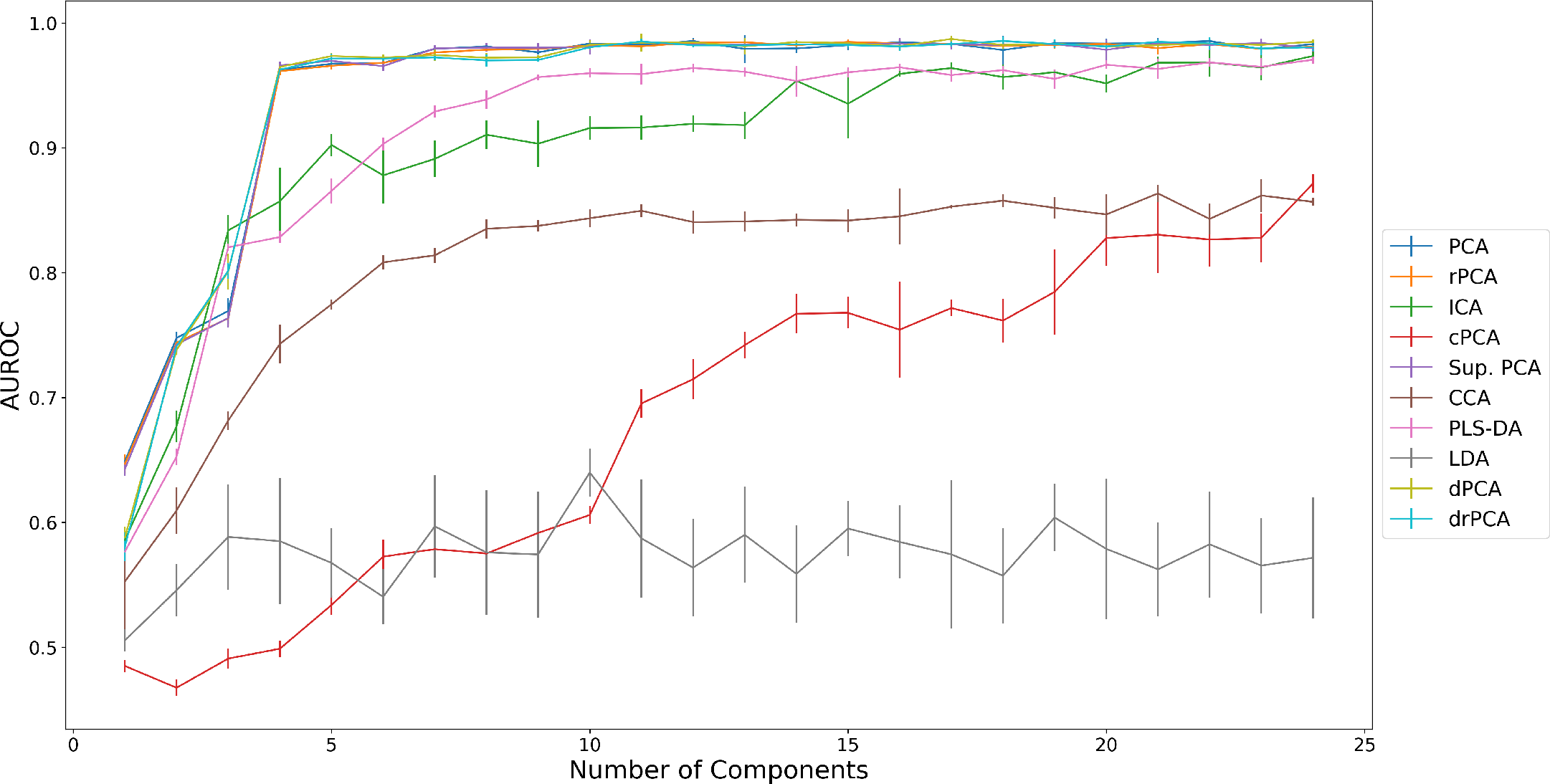
Prediction of tissue of origin from the Combined dataset, with representations learned using case/control labels. The y-axis is the areas under the Receiver Operating Characteristic curves (AUROCs) of the prediction, with x-axis indicating the number of components used in the representations. Errorbars indicate the standard deviation over 3 train/test splits. The representations from LDA, CCA, and PLS-DA underperform most unsupervised methods, demonstrating that the components are not biologically meaningful. In contrast, the drPCA representations are among the best-performing.

From this ranked list, we count the number of selections annotated as oncogenes or tumor suppresor genes (TSG) in COSMIC (24) at each rank. As shown in Figure 6, the differential components give the highest weight to these oncogenic processes.

For a finer-grained anlaysis, we inspect the loadings of the components and variable selection patterns of each of the methods. Shown in Table 1 are the 5 highest weighted genes in the top component for each method; in addition to the methods for components analysis, we also compare to the results of the LIMMA differential enrichment test (25), as compiled by GEPIA (26). Traditional components analyses tend to give higher weight to genes which are associated with high-variance processes, such as cell cycle or ribosomal proteins. In contrast, the differential components select variables more directly related to tumor generation. Each of the top 5 genes selected by drPCA has been implicated as a driver mutation according to DriverDB (23). The components which differentiate tumor samples from control samples are indeed mostly composed of genes known to be associated with cancer.

**Fig. 6.**
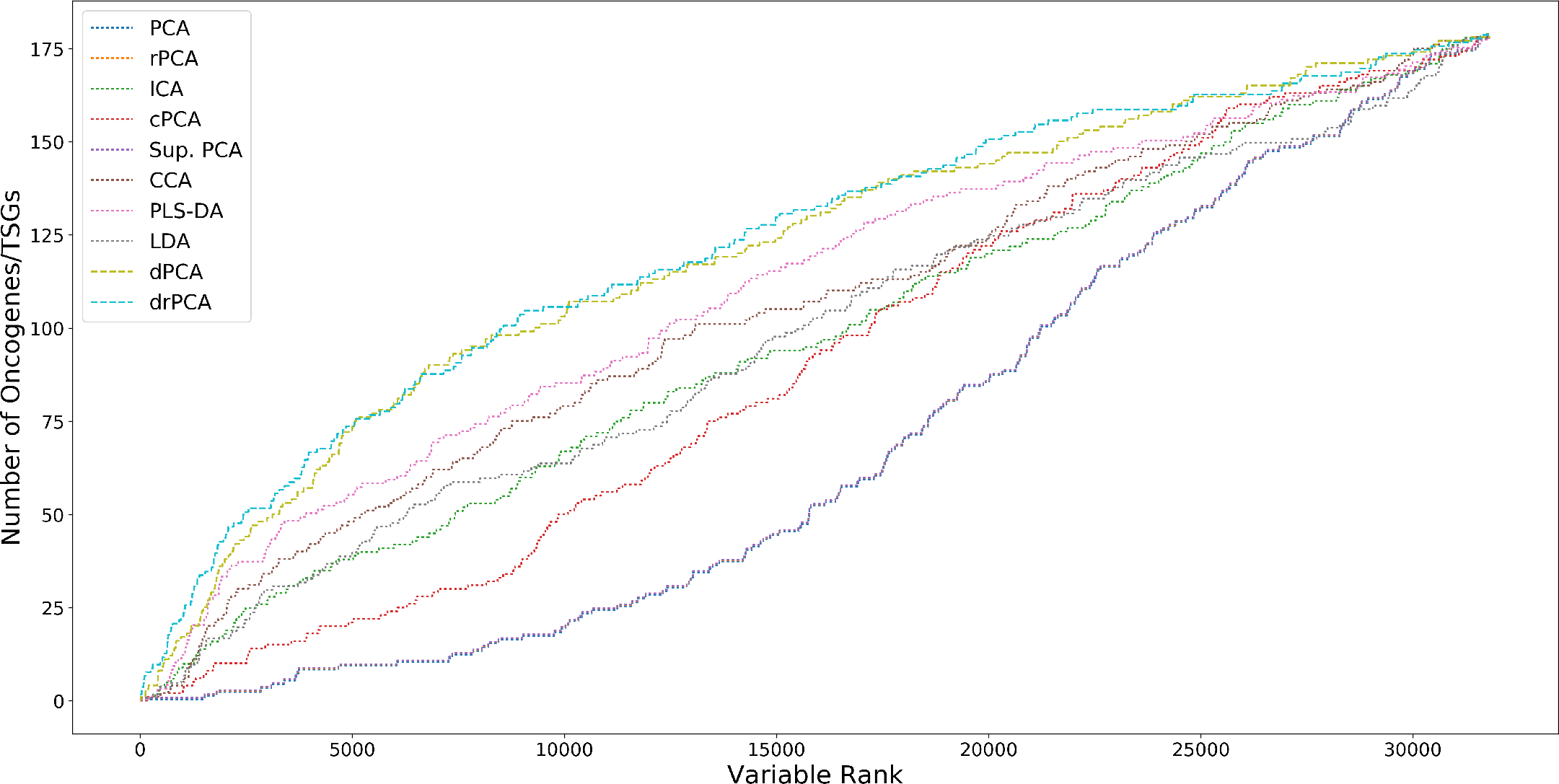
Oncogene/Tumor Suppresor Gene (TSG) selection according to weight in the first component estimated from the Combined cancer dataset. Differential methods give the highest weights to cancer-associated genes.

**Table 1.** The most highly weighted genes in the first c omponent t hat e ach method recovers from the Combined cancer dataset. Genes associated with tumor driver mutations (as annotated in DriverDB(23)) are indicated by a * symbol. The drPCA method gives highest weight to putative driver genes, while many baseline methods pay most attention to high-variance processes, often ribosomal proteins.

| LIMMA | PCA | ICA | cPCA |
| --- | --- | --- | --- |
| KIAA0101* | RP1-102G20.2 | TBC1D9* | MYH1* |
| PAFAH1B3* | RNY1P15 | FOXA1* | ACTC1* |
| C4 | OFD1P17 | RAB30* | AC005616.2 |
| F2R* | RP1-102G20.5 | RP11-102N12.3 | DBET |
| ARHGAP6* | RP1-102G20.4 | DACH1* | MYOG* |
| Sup. PCA | CCA | PLS-DA |  |
| RP1-102G20.2 | VEGFD | AOC3* |  |
| RNY1P15 | RP11-257I8.2 | RP11-736K20.4 |  |
| OFD1P17 | CLEC3B* | VEGFD |  |
| RP1-102G20.5 | RP11-193H18.3 | CLEC3B* |  |
| RP1-102G20.4 | CA4* | NPR1* |  |
| LDA | dPCA | drPCA |  |
| GLYAT* | ZSCAN25* | ZSCAN25* |  |
| CIDEC* | C2orf42* | C2orf42* |  |
| ADIPOQ* | FBXO42* | FBXO42* |  |
| TRHDE-AS1 | FAM133B* | FAM133B* |  |
| C14orf180* | C2orf49* | RBM33* |  |

## Discussion

Summarizing the ways in which samples differ between experimental conditions is a central task in scientific inquiry, but made difficult by the large dimensionality of modern bioinformatics datasets. Common methods of unsupervised dimensionality reduction produce components which do not distinguish between experimental groups. This problem is worsened by the selective pressure of the observations in gene expression studies; axes of major variation in the data often correspond to unregulated, and largely noncritical, processes. In this paper, we have presented a way to extract the processes which differentiate experimental conditions by adapting unsupervised techniques to the supervised setting. Our framework can outperform methods designed specifically for supervised data when denoising methods, such as in drPCA, are used.

We are interested to see the variety of unsupervised dimensionality reduction techniques that can be repurposed in this supervised setting. For instance, we may want to estimate components which correspond to genetic pathways, similarly to (27, 28). This can be accomplished under the differential PCA framework by using a Bayesian PCA method with a prior that links genes according to pathway annotations. Even without these additional biological information, we have shown that drPCA outperforms common supervised dimensionality reduction methods at producing biologically-meaningful components.

## Conclusions

In this paper, we have considered whether components analysis can compress samples from case/control studies onto biologically meaningful axes. We found that PCA and unsupervised variants do not separate case and control samples due to overemphasis on high-variance processes such as cell cycle and ribosomal processes. Supervised methods of dimensionality reduction separate case and control samples, but the resulting components have questionable biological utility and are overfit to sample labels. To address this problem, we have presented differential PCA: a method which applies PCA on the set of difference vectors between the samples. Our new method Differential Robust PCA (drPCA) effectively identifies axes of differentiation, outperforming standard supervised methods such as PLS-DA even under noisy conditions. When applied to gene expression data of cancer patients, drPCA produces components which summarize oncogenic processes. In future work, we are interested to incorporate prior biological knowledge to extract pathway-level components.

## Acknowledgements

Thanks to Bryon Aragam, Haohan Wang, Michael Kleyman, and Ziv Bar-Joseph for helpful discussion about these ideas.

## Funding

This work is supported by the National Institutes of Health grants R01-GM093156 and P30-DA035778.

## Footnotes

1 www.Github.Com/Blengerich/Drpca

2 cancergenome.nih.gov

